# Roles of dimeric intermediates in RNA-catalyzed rolling circle synthesis

**DOI:** 10.1101/2024.05.14.594117

**Authors:** Emil Laust Kristoffersen, Ewan K. McRae, Niels R. Sørensen, Philipp Holliger, Ebbe S. Andersen

**Author notes:** contributed equally to the work (Emil Laust Kristoffersen, Ewan K. McRae). jointly supervised the work (Philipp Holliger, Ebbe S. Andersen). Communication to & (Philipp Holliger, Ebbe S. Andersen).

## Abstract

The RNA world hypothesis is supported by the discovery of RNA polymerase ribozymes that can perform RNA-catalyzed RNA replication processes on different RNA templates. Recently, RNA-catalyzed rolling circle synthesis (RCS) on small circular RNA (scRNA) templates has been demonstrated. However, the structural and dynamic properties of scRNA replication and its products and intermediates have never been explored. Here we have used cryogenic electron microscopy (cryo-EM) to characterize products and intermediates relevant for RCS replication and find that these form an unexpectedly diverse group of RNA nanostructures. The main structural motif observed is a fully hybridized dimeric complex composed of two scRNAs and their complement strands resolved to 5.3 Å. Cryo-EM also reveals higher order dimer filaments and dimer assembly intermediates suggesting a mechanism for assembly of the observed complexes. We show that the dimer complexes are stable and inhibit RNA-catalyzed RCS, but can be reactivated by addition of more scRNA templates. We propose that dimer formation may be a general property of RCS replication and speculate that the observed dimers might have benefited a primordial RNA genetic system by providing a stable “storage” form of RNA replication products and by coordinating RNA replication on both scRNA template strands.

## Introduction

How life and its first genetic system may have emerged on the early earth is an area of active investigation. Among various competing ideas, the so-called RNA world hypothesis^1,2^ remains an attractive proposition due its conceptual simplicity and the compelling (if circumstantial) evidence in modern biology. This includes the role of RNA catalysts (ribozymes) in universal biological processes such as translation and RNA processing, and the presence of ribonucleotide cofactors in central metabolic processes. The RNA world is further supported by recent significant progress in the demonstration of increasingly plausible prebiotic chemistry routes towards the synthesis of activated ribonucleotides^3–7^ and the non-enzymatic, templated polymerization of RNA^8–11^. While a cornerstone of the RNA world hypothesis - RNA-catalyzed RNA self-replication - remains to be demonstrated, *in vitro* evolution has enabled the discovery of RNA polymerase ribozymes with ever increasing synthetic power^12^ including the synthesis of active ribozymes^13,14^ and longer-than-self RNA products on some templates^15^ and the processive polymerization and promotor recognition^16^. Recently, a polymerase ribozyme, that utilizes tri-nucleotide triphosphates (pppNNN, triplets) rather than ribonucleotide triphosphates (NTPs) as substrates, has been described^17^. This triplet polymerase ribozyme (TPR) is a heterodimer composed of a catalytic subunit (5TU) and an inactive accessory subunit (t1) as confirmed by its cryogenic electron microscopy (cryo-EM) structure ^18^. This TPR has been shown to be able to copy highly structured templates as well as segments of itself^17^.

While there is thus significant progress in obtaining increasingly active polymerase ribozymes, RNA replication will likely also require a strategy for strand separation as duplex RNAs are exceedingly stable. Among different, conceivable strategies, rolling circle synthesis (RCS) is an attractive mode as it mimics the simple replication strategies of viroids (and some viruses) in biology, which have themselves been suggested to predate cellular life^19^ and be relics from the RNA world^20^. However, these natural replicons use protein-based polymerases to replicate, which were likely not present in an RNA world. An important feature of RCS is that the energy released from the polymerization reaction can - in principle - be directed to drive strand displacement^21,22^ and result in cotranscriptional displacement and folding of the nascent strand. Therefore, RCS potentially enables linkage of genotype (RNA template) with the RNA phenotype (folded nascent strand) - a prerequisite for evolution - without compartmentalization. In contrast, polymerization on linear strands leads to formation of highly stable RNA duplexes that require an external driving force such as temperature, wet-dry-, or pH-cycling for continued replication and to assemble an active state^23,24^. While RCS requires the formation of circular RNA (circRNA) templates, these have be shown to arise from self-ligation processes^25,26^, or wet / dry cycling^27,28^ suggesting several prebiotic routes to circRNAs.

Recently, we reported that the TPR can perform RCS on small circular (sc)RNA templates^29^. The RCS was made possible due to the fact that hybridization of small RNA circular templates and their growing nascent strand is energetically disfavored, if the scRNA is considerably smaller than the duplex RNA persistence length of approximately 300 base pairs (bp)^30^. Indeed, molecular dynamics (MD) simulation showed that a circular template of 36 nt could maximally form a 24-bp duplex by hybridizing with its complementary linear strand. Upon further extension (beyond 24 nt), the MD simulation suggested that the system is dynamic, leading to unpaired 5’- and 3’-ends of the nascent strand and unpaired segments in the scRNA template, which we proposed to facilitate displacement of the 5’-end from the template upon iterative extension of the 3’-end. However, while the TPR was able to synthesize nearly three rounds on the scRNA template^29^, we also observed a strong inhibition of synthesis after completion of just one round on the scRNA template, which was challenging to rationalize.

To gain deeper insight into the RCS process and a better understanding of the observed strong inhibition upon full length circle synthesis, we used cryo-EM to study the complex formed by the full-length circle (scRNA) and nascent (complementary) strand (cmpRNA). Unexpectedly, cryo-EM revealed a diverse variety of distinct structural species identified as monomers, dimers and multimers of the scRNA-cmpRNA complex. The main structural complex, that was resolved to a resolution of 5.3 Å, was composed of two scRNAs and two cmpRNAs forming two interconnected parallel RNA helices. This dimer structure was further investigated in primer extension assays confirming it as an inhibitory product that could however be reinitiated by the addition of an excess of scRNAs.

## Results and Discussion

### Cryo-EM structures of the full-length RCS product

We previously demonstrated RNA-catalyzed rolling circle synthesis (RCS) through primer extension of a nascent strand on a scRNA template by a triplet polymerase ribozyme (TPR) (Figure 1A)^29^. For our analysis, we focused on the 1:1 complex formed between a full-length 36-nt nascent strand (cmpRNA) and a 36-nt circular template (scRNA), since this represents a previously observed point of strong inhibition of the RCS reaction^29^. Denaturing gel electrophoresis showed that cmpRNA and scRNA have distinct electrophoretic mobilities (Figure 1B, lane 1 and 2). However, when combined, they form a distinct band with lower mobility (Figure 1B, band II in lane 3), which is stable under denaturing conditions suggesting formation of a highly stable complex. Next, we performed native gel electrophoresis to verify the formation of a complex between circular and linear RNA, using radiolabeled cmpRNA and unlabeled scRNA (Figure 1C). In addition to a major band (Figure 1C, band II) that may correspond to band II in Figure 1B we observed two slower migrating bands (band III and IV) that are not present in Figure 1B and may therefore represent weaker interacting multimers of band II.

**Figure 1.**
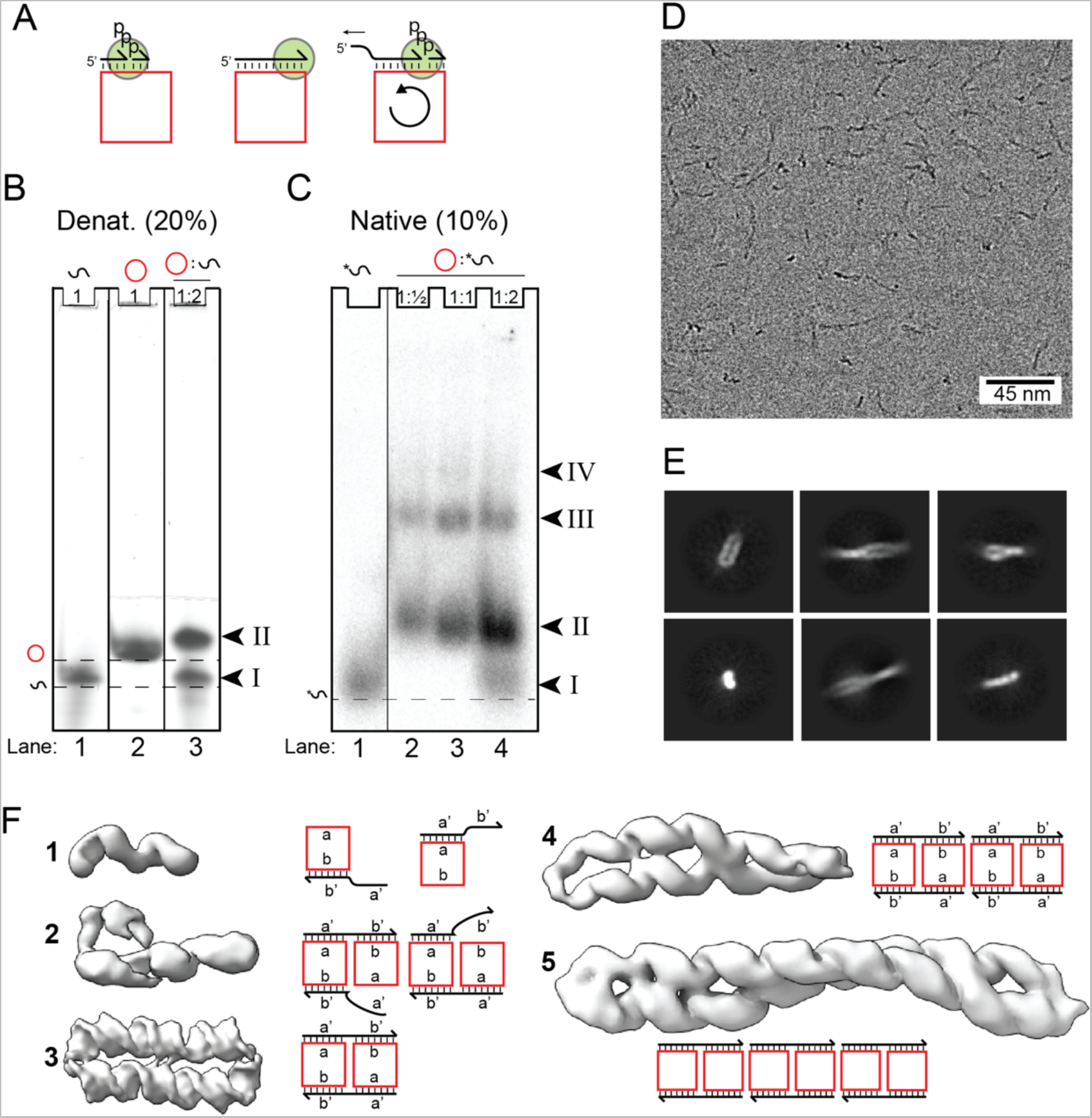
Complexes between circular template and linear product strands. (A) Schematics showing the basics of RCS, where the scRNA (red) binds a primer (black) that is elongated at the 3’-end (arrow) by an RNA polymerase ribozyme (green). When RCS reaches a certain length, the 5’-end is released from the template due to strain in the scRNA. (B) 8M Urea denaturing (20%, 19:1) polyacrylamide gel with RNA constructs. RNA was stained with SybrGold. Arrowheads and roman numbers denote bands in the gel. The dashed line represents the front of band migration of selected species. (C) Native (10%, 37:1) polyacrylamide gel. Bands observed in the native gel represent 5’-end radiolabeled cmpRNA. Whole raw images of gels shown in (B) and (C) are found in Figure S1 (A) and (B), respectively. (D) A representative cryo-EM micrograph. (E) Selected 2D classes. (F) The five structural classes observed by cryo-EM and their interpretations as monomer, dimer, and multimer species. Domains of the circle are denoted a and b and the complementary domains on the product strand are denoted a’ and b’.

Next, a sample with equimolar amounts of scRNA and cmpRNA was analyzed by cryo-EM. The micrographs and class averages revealed both distinct particles and longer filaments (Figure 1D, Figure S2-6). Using an iterative 3D reconstruction approach, we identified 5 distinct structural classes (Figure 1E, Figure S7-11, for details see Materials and Methods). All classes showed clear helical features, providing confidence in identifying them as genuine RNA complexes. Class 1 showed a density corresponding to two helical turns of an RNA double helix. Class 2 showed density similar to two parallel RNA helices, one long and one shorter. Class 3 led to our best resolved structure showing two parallel helixes each having slightly more than 3 helical turns. We presume that this structure corresponds to band II in the denaturing gel in Figure 1B as this structure is likely highly stable. Finally, class 4 and 5 displayed long twisted structures approximately double and triple the length of class 3, and likely corresponding to bands III and IV in the native gel in Figure 1C.

Based on the identified structures we concluded that class 3 likely represents a homodimer between two scRNA-cmpRNA complexes that have combined by mutual cross-hybridization driven by the geometrical restriction of the small circular template and the rigidity of the RNA duplex. Class 4 and 5 represent multimers formed by end-to-end stacking of the class 3 dimers. Class 1 and 2 were poorly resolved but resemble RCS structures as predicted by MD simulation and may represent intermediates trapped on the way towards formation of the dimer. Furthermore, class 2 has the potential to form via two different pathways where either the 5’- or 3’-end does not anneal. These interpretations are schematically illustrated next to the density maps in Figure 1E. Next, we examined the individual structural classes in more detail, starting with class 3.

### Structure of fully hybridized dimer

The dimer from class 3 was found to be composed of two scRNAs and two cmpRNAs that are fully hybridized (Figure 2A). The cryo-EM reconstruction using C2 symmetry led to our highest resolution map of 5.3 Å that revealed two parallel RNA helices of 36 bps connected by antiparallel crossovers formed by the two scRNAs – a central two-strand crossover and two proximal one-strand crossovers (Figure 2B and Figure S9 for comparing C1- and C2-symmetry). A non-ideal crossover spacing is observed, with 22 bp on one helix and 14 bp on the other. The 22-bp segment has two helical turns between the crossovers (corresponding to an ideal spacing of 11 bps per turn), whereas the 14-bp segment only has 1 turn. The non-ideal crossover length of the 14-bp segment results in its underwinding and a global distortion of the structure where the 22-bp helix bends outwards and the 14-bp segment bends inwards giving the structure a slightly curved shape (Figure 2B).

**Figure 2.**
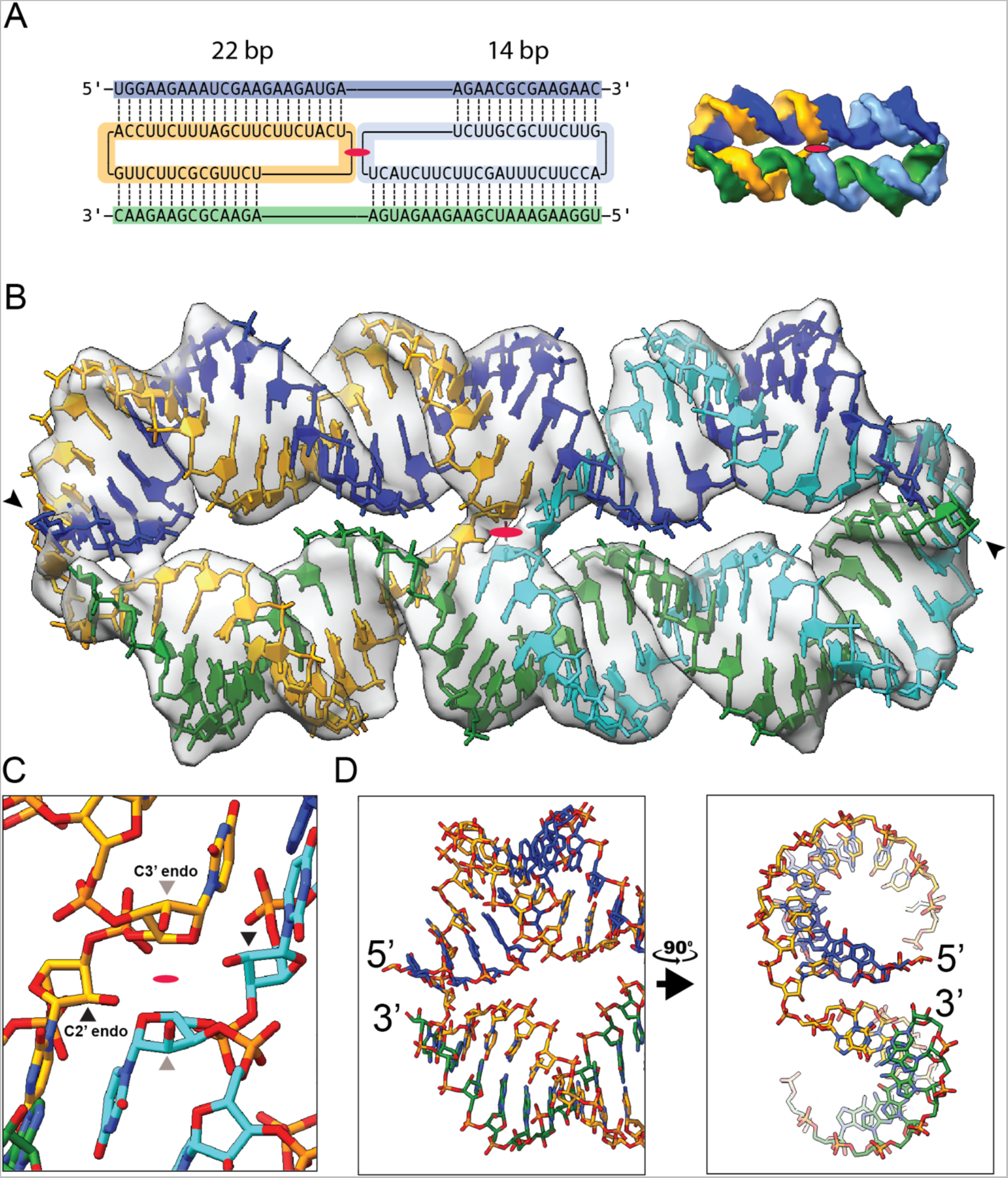
Structure of fully hybridized dimeric circle-product complex. (A) Secondary and tertiary structure model of two scRNAs (orange and cyan) bound to their product strands (blue and green) with domains of 22 and 14 bps as suggested from the cryo-EM data. Red ellipse indicates C2 symmetry axis. (B) Atomic model based on the density of cryo-EM class 3. Colors as in panel A. Arrows indicate 5’-end. Red ellipse indicates C2 symmetry axis. (C) Zoom on crossover junction where the 5’ nucleotides entering the junctions are found in the C2’-endo conformation. (D) Zoom on helix ends where the scRNA crossover from one helix to the other is shown in two perpendicular views.

Investigation of the central crossover in the dimer shows that it adopts a similar conformation as has been observed in cryo-EM structure of crossovers in RNA origami^31^, where a C2’ endo conformation is adopted by the nucleotide at the 5’-end of the strand entering the crossover, whereas the 3’-end of the strand entering the crossover is in the C3’-endo conformation (Figure 2C). The C2’ endo conformation allows the 3’-end of the nucleotide to point out of the helix to facilitate formation of the crossover. The one-strand crossovers connecting the helices at the two ends of the complex are formed by nucleotides with C3’-endo conformation (Figure 2D). The distortion of the helices results in a proximity of the 5’- and 3’-ends of the two different cmpRNAs, where the 5’-end of one cmpRNA projects above the 3’-end of the other cmpRNA.

### Twisting filaments produced by stacking of dimers

The density maps for class 4 and 5 could be readily modelled by stacking class 3 dimers to form twisted filaments – with class 4 comprising two dimers and class 5 three dimers (Figure 3A and B). These multimer models suggest that the interaction between dimer units in the filament are mediated by base-stacking of terminal bases from each unit (Figure 3C and D). Based on the models we suggest that A20 and G21 of one circular strand stacked with G21 and A20 of the other circular strand, respectively, and the 5’-end U1 of one unit stack on the U1 of the second unit, while the 3’-ends were non-stacking. The 5’-ends point outwards from the plane of the interaction and 3’-ends were point inwards. The described distortion in the complex caused a global twist in the filament (period of ∼4 dimer units/turn) (Figure 3B).

**Figure 3.**
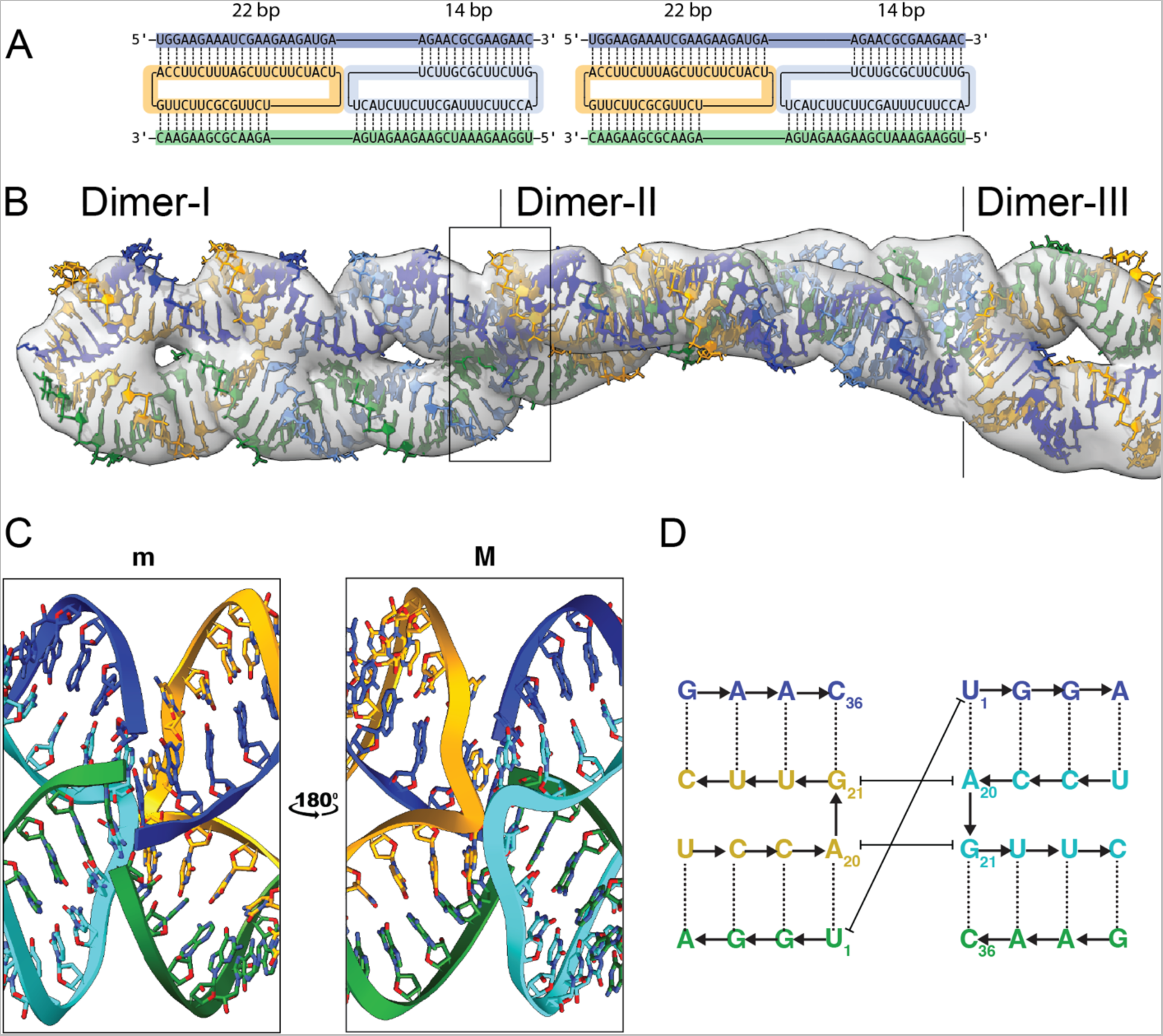
Multimeric structures composed of stacking homodimers. (A) Secondary structure of two homodimers attached end-to-end. Red ellipse indicates C2 symmetry axis. (B) Cryo-EM map of class 5 with superimposed atomic models. Colors as in panel A. (C) Zoom on stacking of the end-to-end crossover junction shown perpendicular to the C2 symmetry axis from the minor (m) and major (M) groove side. (D) Secondary structure diagram showing the base stacking interactions between the terminal bases.

Notably, the 22-14 segmentation of the dimer also leads to a misalignment of the 5’- and 3’-ends of adjacent units. Indeed, this can be tested by ligation. Our structural model predicts that the dimers would ligate poorly. In contrast, scRNAs with a circumference corresponding a whole integer of the RNA helical pitch (33 nt), would - despite also being distorted – likely present 5’- and 3’-ends better aligned for ligation. We tested this hypothesis using ligation experiments with dimers prepared from scRNAs and corresponding cmpRNAs that were either 33 nt or 36 nt in size (Figure S12 and Table S1). We observed a more intense low mobility ligation band for the 33 nt circle compared to the 36 nt circle, supporting our observed misalignment of the 5’- and 3’-ends in class 4 and 5 models.

### Partially hybridized structures

The low resolution cryo-EM maps of class 1 and 2 are consistent with partly hybridized scRNA and cmpRNA. The density map for class 1 was observed to have slightly more than two helical turns. The density was analyzed by aligning a model of the scRNA-cmpRNA complex that was previously predicted by MD simulation to form approximately 24 bps^29^ (Figure S13). The similarity between the class 1 map and the MD model indicates that they represent the same species. Based on this we constructed a new 3D model of the whole system that was fitted to the class 1 density (Figure 4A and B). Note that the cryo-EM density shows only the double-stranded segment since the single-stranded regions are flexible and are averaged out during 3D reconstruction. In Figure 4A we show the most G / C rich 24 bp stretch since this will be the most thermodynamically stable structure.

**Figure 4.**
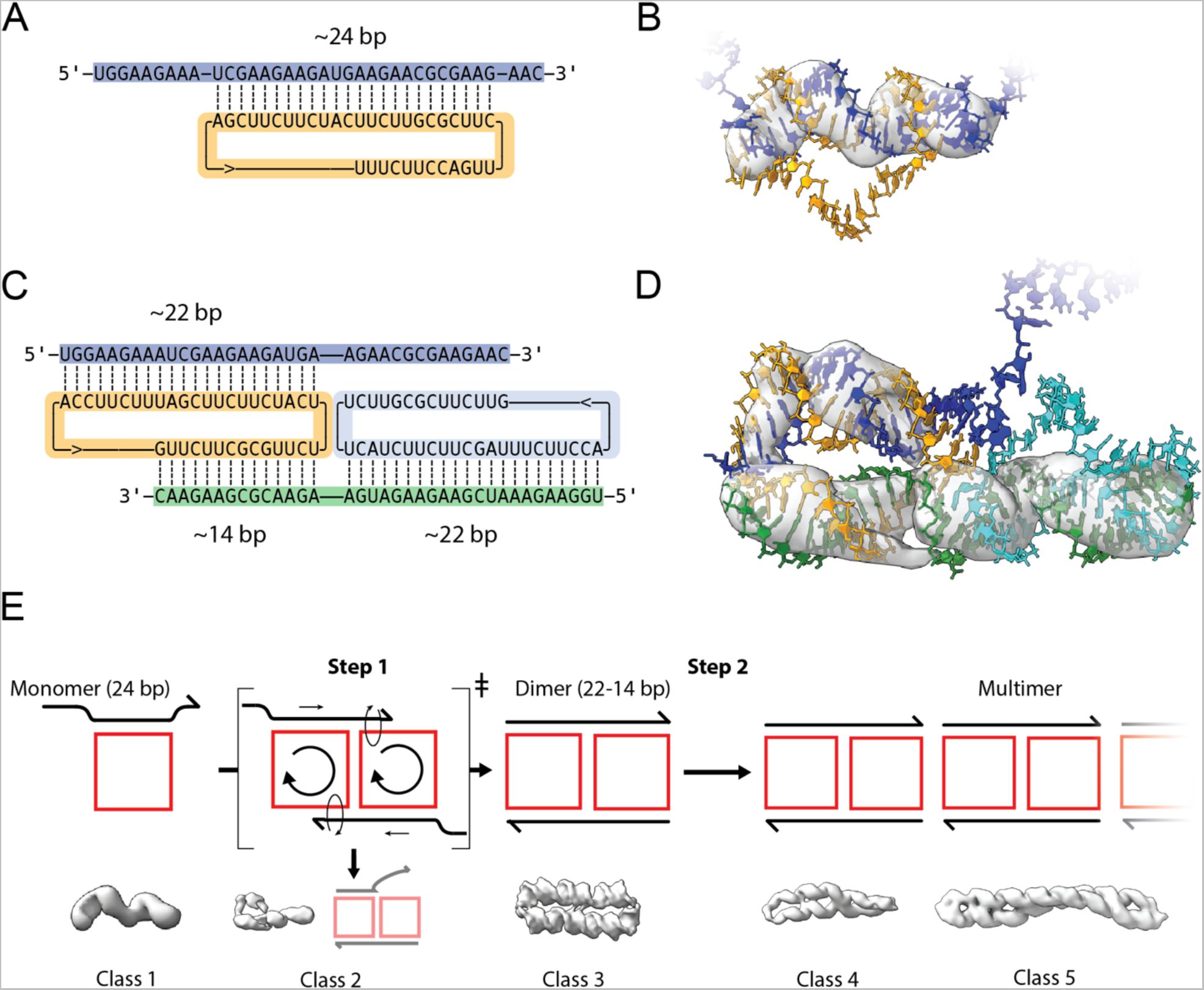
Partially hybridized complexes and hypothetical self-assembly path. (A) Secondary structure model of the scRNA (orange) bound to its product strand (blue) by ∼24 bp. (B) Density of cryo-EM class 1 superimposed on atomic model. (C) Secondary structure model of two not fully hybridized scRNAs (orange and cyan) bound via their product strands (blue and green). (D) Density of cryo-EM class 2 superimposed on atomic model. (E) Illustration and density maps of the proposed mechanism for scRNA and cmpRNA monomers to dimerize via a rolling circle mechanism transition state leading to formation of the dimer (class 3) (Step 1). Step 2 leads to multimerization (class 4 and 5) by stacking dimers. Stalled folding intermediates of class 1 and 2 is proposed to emerge as retained monomers or as a failed dimerization intermediate as illustrated under the respective complexes. The proposed assembly path is shown in Movie 1.

The density map for class 2 showed low-resolution densities consistent with models composed of two scRNAs and two cmpRNAs, where the second cmpRNA had only partly hybridized. Based on the dimer model derived from class 3, we propose a model for class 2, where one cmpRNA is fully hybridized and the other cmpRNA is partly hybridized (Figure 4C and D). Based on the position of the major grooves in the class 2 map, we can deduce that it is the 3’-end of one of the cmpRNAs that does not hybridize (as seen in Figure 4C and D).

### Proposed self-assembly pathway for dimers and multimer

The observed structural classes are related and seem to be on an increasingly thermodynamically stable folding path. In class 1 only ∼66% of the possible base pairs (bp) (∼24 out of 36 bp) are formed, in class 2 ∼80% (∼58 out of 72 bp (36 bp per circle)), and in class 3 100% of the possible bp (72 out of 72 bp possible) are formed. Based on this we propose a rolling circle-based assembly pathway to explain the formation of the observed species (Figure 4E). In step 1, two class 1 monomers dimerize and maximize base pairing by a rolling mechanism weave the strands together to form class 3. Rolling of the central crossover is known to occur in mobile Holiday junctions found in DNA recombination intermediates ^32,33^ and has formerly been utilized in DNA nanotechnology ^34,35^. Interestingly, we observe a trapped state of the rolling mechanism, where only one of the unpaired ends have hybridized (Figure 4E, class 2). In step 2, class 3 dimers assemble into multimers by end-to-end stacking, leading to the filament-like class 4 and 5 structures. Our proposed assembly pathway is illustrated in Movie 1.

### The dimer as a roadblock and substrate for RCS

The structural study of scRNA and cmpRNA complexes revealed an unexpected diversity of structures that represent likely intermediates of RNA-catalyzed RCS as the nascent strand is extended towards full length (Figure 5A, step 1-2). One prediction from our studies would be that the class 3 dimer would be a dead-end product and inhibit further 3’-end extension in the RCS reaction both due to its stability and poor accessibility of 3’ ends. To test this, we mixed equimolar scRNA and cmpRNA and investigated if the cmpRNA 3’-end in a gel purified class 3 dimer could be extended by the TPR. As predicted, our data showed that the cmpRNA could not be extended by the TPR once incorporated into the dimer (lane 2 in Figure 5B and Figure S14) even upon prolonged incubation. This observation now allows a rationalization of the previously unexplained observation of strong RCS inhibition at one full scRNA length extension (as observed in ref.: ^29^). Furthermore, it is consistent with the observation that dilution decreased this inhibitory effect (also observed in ref.: ^29^), as dilution may slow dimer formation resulting in more class 1 monomer structures compatible with RCS.

**Figure 5.**
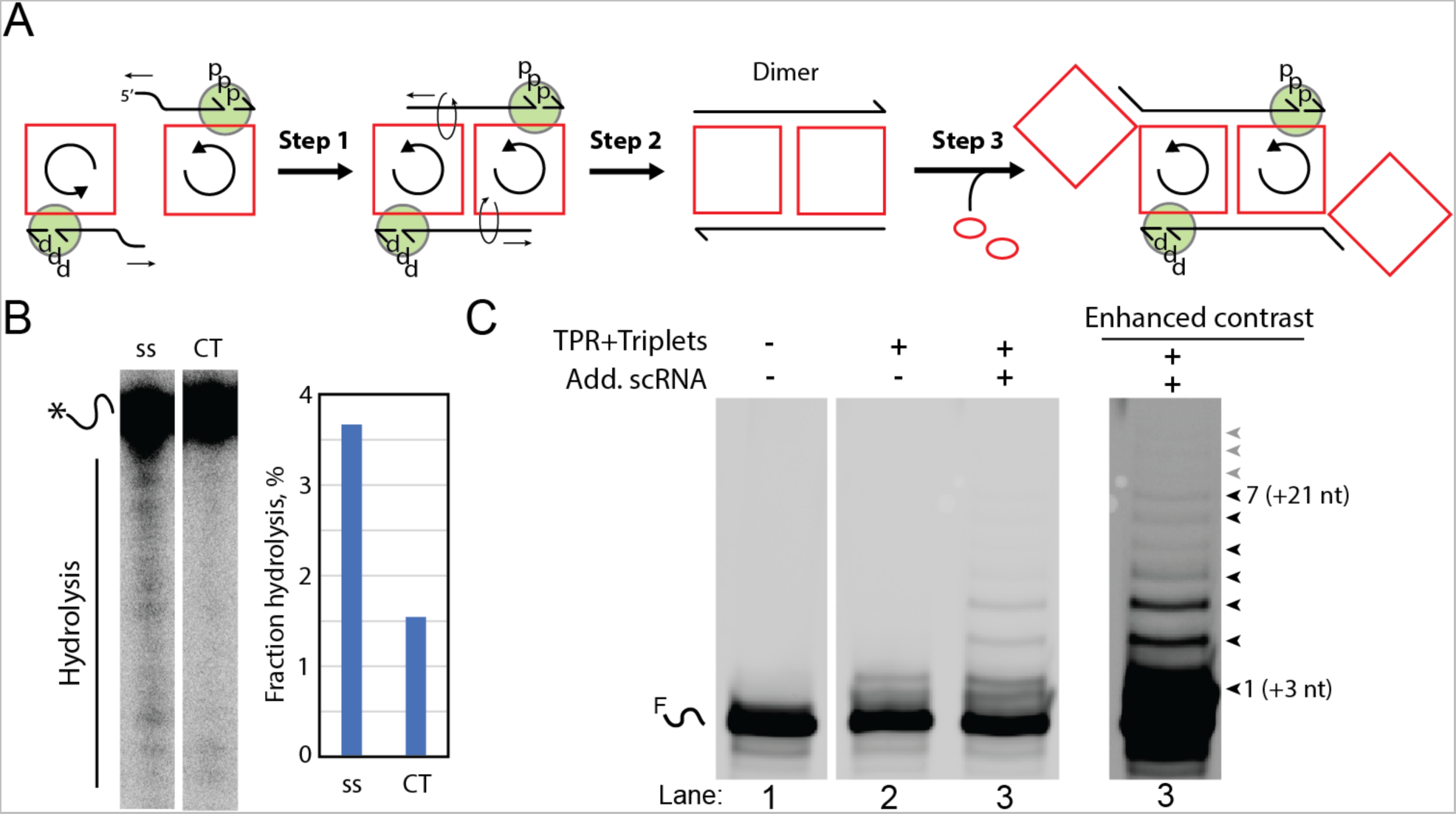
Dual rolling circle replication. (A) Illustration of how two rolling circle replication complexes can dimerize by annealing of the displaced product strand to the second scRNA (step 1). Step 2 illustrates how continuous extension of the product strand allows it to gradually weave the scRNAs together finally forming the dimer. Finally, step 3 illustrates the continued synthesis (see Movie 2). The green circle symbolizes the TPR. (B) Denaturing gel (10% 19:1) show hot labeled cmpRNA hydrolysis after incubation (1 week at -7 degrees Celsius) under reaction conditions. The cmpRNA was either incubated as ssRNA or as part of the dimer. Barchart of quantified hydrolysis shows that the cmpRNA was less stable as ssRNA compared to when it was in the dimer. (C) Denaturing gel (20% 19:1) show experiments where dimer was incubated with active TPR to investigate if this would lead to extension of the cmpRNA. Lane 1 shows fluorophore labelled cmpRNA alone as a control. Lane 2 shows results from an experiment where equimolar cmpRNA and scRNA were preincubated to form the dimer prior to incubation with the active reaction conditions (TPR + triplets). Lane 3 shows results from a similar experiment but where additional scRNA was added to preincubated dimer and then incubation with active reaction conditions. Elongation of the cmpRNA was only observed in lane 3. Same gel is also shown with enhanced contrast to underline the extend of the cmpRNA elongation. Note that 10-fold excess of unlabeled cmpRNA was added to the samples used in (B) and (C) (and in Figure S16) before denaturing gel analysis to allow release of the of the labeled cmpRNA from the dimer, (see Material and methods section). Whole raw images of gels shown in (B) and (C) are found in Figure S1 (C) and (D), respectively.

Next, we wondered if the unusual stability of the dimer would not only prevent further RCS replication but also attenuate degradation. Indeed, when we investigated the dimer stability, we found that cmpRNA was protected from hydrolysis being over 3-fold stabilized when it was part of the dimeric circle / template (CT) complex compared to when it existed as single strand (ss) alone (Figure 5B).

The rolling circle hybridization mechanism described above suggests that the class 3 dimer might replicate via a coordinated “two-fold RCS” by a rolling motion of both scRNA templates that would allow the annealing and iterative addition of new triplets by TPR on both strands (see proposed mechanism in Movie 2). Inspired by controlled rolling motion in a DNA actuator^34,35^, we hypothesized that the replication block might be relieved by addition of additional scRNA templates to allow both cmpRNAs of the dimer to slide and facilitate additional templated elongation of the 3’-end (Figure 5A, step 3). This did indeed restart the extension reaction from previously inert dimer and multiple triplets were added to the cmpRNA strand even though with modest efficiency (Figure 5C, lane 3).

### Hypothesis for role of dimer in replication

The above experiment shows that while formation of the stable class 3 dimer inhibits RCS, inhibition can be relieved (in part) by the addition of more scRNA templates. We propose that this continued extension of the 3’-end is facilitated by the hybridization of the 5’-ends to the scRNAs and the triplet substrate to the scRNA template at the 3’-end of the cmpRNA (Figure S15A). A similar mechanism might also be driven by internal folding of the 5’ region of the nascent strand preventing reannealing to scRNA (Figure S15B). As suggested by the structure of the dimer, the 5’-end of one cmpRNA projects above the 3’-end of the second cmpRNA (Figure 2D), thus priming the complex to roll in the correct 3’-5’ direction and inhibit rolling in the reverse (5’-3’) direction. Reverse rolling is also prevented by the ratchet-like RCS reaction with triplet addition to the 3’ end - an essentially irreversible process.

The dimer has intriguing properties both as an RNA storage form (dead-end complex) and as a scaffold for coordinated two-fold RCS: Firstly, the dimer is fully hybridized thus having increased stability compared to the monomer (Figure 5C). Secondly, the nascent strand may fold into an active conformation (e.g., an RNA polymerase ribozyme) that has the potential to act on itself (in cis) within the replication complex (Figure S15B). For the monomer RCS the distance between the 5’-end and the position of the ligation is 8 nm for a 36-nt template and increases with template length. However, for the dimer RCS the distance between the 5’-end and the nearest ligation site is only 4 nm and does not increase with template length. Thirdly, the rolling mechanism of the dimer might function as a type of proof-reading mechanism for mis-incorporated bases (including bases with non-cognate chemistry or chirality) (Figure S15C). As the rolling relies on the isoenergetic relation between 3’-end base pair formation and base pair melting in the 5’-end, incorporation of mismatched bases at the 3’ end will destabilize 3’-end hybridization and in turn disfavor coordinated 5’-end melting and thus inhibit extension as well as potentially expose mismatched, single-stranded 3’-ends to increased hydrolysis. If working, this mechanism would make sure that only “proof-read” sequences would make it through the cogwheels of the structure and get protruded in the 5’-end. Finally, because this RCS would be a coordinated process, misincorporation on one scRNA would also inhibit further extension on the second scRNA increasing fidelity of replication on both templates simultaneously.

When docking the structure of two ribozymes (the class I ligase ribozyme (cIL) (PDB:3IVK) and TPR, (PDB: 8T2P)) onto the structure of the model of class 3 dimer so the active sites of the ribozymes were positioned on the dimer at the 5’-end of an annealed triplet (Figure S15D and E for cIL and TPR, respectively) it becomes clear that cIL could in theory access the ligation junction (Figure S15D), while TPR was hindered from binding to the two ends of the dimer simultaneously due to steric clash of the noncatalytic subunits (t1 in Figure S15E). The steric clash is a likely part of the explanation for the low efficiency of the extension on the dimer by the TPR but might be solved in the future through new and improved RNA polymerase ribozymes selected for circular RNA templates. Indeed, cooperative 3’-end extension by two ribozymes might provide sufficient energetic driving force for double rolling scRNA mechanism without the addition of additional scRNA templates for activation.

In this study we have investigated the properties of a 36 nt circular template but many different sizes of RNA circles could conceivably be investigated for their properties in forming the dimer and their function as a template for two-fold RCS. Our prediction would be that scRNAs in the length range of 22 to several hundred bps would likely at least to some extent form dimer-type structures due to the persistence length of the double-stranded RNA helix of approximately 300 bps^30^. We also show that these dimers must either be avoided or reactivated to get an efficient RCS. Designing new experiments taking these factors into consideration as well as designing optimized ribozymes might lead to more efficient RCS.

## Conclusion

In this study we have used cryo-EM to investigate the RCS process on scRNA templates by the structural analysis of putative RCS intermediates and products. We find that scRNAs and cmpRNAs of 36 nt in length can form of an unanticipated diversity of structures with different stoichiometries, including a fully annealed dimer (comprising 2 scRNAs and 2 cmpRNAs) and multimers hereof. These complexes form because of the restrictions on hybridization of scRNA templates and the cmpRNAs, which arises due to geometric strain in the scRNA when the circle size is considerably smaller than persistence length of duplex RNA. Furthermore, we show that the dimer structure is a highly stable dead-end complex that inhibits the primer extension by TPR, but inhibition can be relieved by addition of additional scRNAs. We propose a new double circle RCS mechanism based on the dimer structure and its ability to be extended by TPR in the presence of additional scRNA templates. Our study further underlines the importance of the template structure and topology in RNA-catalyzed replication and outlines how rolling circle replication mechanisms might have provided fitness benefits in prebiotic RNA replication.

## Material and methods

### RNA preparation

RNA was prepared essentially as described in ref.: ^29^. In brief, RNA was *in vitro* transcribed or chemically synthesized (IDT). When relevant, polynucleotide kinase (NEB) was used to remove 2’-3’ cyclic phosphate. scRNA was prepared by T4 RNA ligase 1 ligation (NEB) exactly as described in ref.: ^29^. RNA was then gel purified by 20% (denaturing) urea PAGE, detected by UV-shadowing, and recovered by freeze and squeeze extraction (removing gel pieces with a Spin-X column (0.22 μm pore size, Costar)). When relevant cmpRNA (500 pmol) was 5’-end phosphate-32 “hot” labeled using fresh gamma phosphate-32 labeled ATP (50 nmol) and PNK (NEB) following the producer’s recommendations. Excess gamma phosphate-32 labeled ATP was removed during PAGE purification. Fluorophore-labelled cmpRNA was commercially acquired (IDT). Eventually this procedure led to gel-purified circularized scRNA and gel-purified cmpRNA (in some cases labeled with radioactive 5’-phosphate or FAM fluorophore).

### RNA circularization

Gel purified linear scRNAs were circularized as described in ref.: ^29^. In short 5’-phosphorylated and 3’-OH RNA was circularized with RNA ligase 1 (NEB) followed by denaturing PAGE purification as described above. The gel band representing circularized RNA monomer (scRNA) was identified as described in ref.: ^29^.

### Assembly of the cmpRNA and scRNA structures and PAGE analysis

Gel-purified cmpRNA (with or without label as specified in the text) and scRNA were mixed in the reaction buffer (50 mM Tris, pH 8, 100 mM MgCl_2_) at noted molar ratios and incubated for 10 min at room temperature and then adjusted to reaction conditions. After this, the samples were stored on ice for up to a few hours until used. Assembled RNAs, including relevant control samples, were analyzed by denaturing (8 M urea) PAGE gel electrophoresis (10% or 20% as noted in the text) running in 1x TBE buffer at 25 W for 1-2 hours or by (native) PAGE gel electrophoresis where the gel was cast with 1xTBE and 100 mM MgCl_2_ and run at 100 V for 16 hours. Gel-separated radioactive RNA was detected using radioactive films (GE HealthCare) and scanned using a Typhoon scanner (Thermo Scientific). FAM fluorophore-labeled RNA was analyzed directly in the gel by a Typhoon scanner.

### Cryo-EM

RNA for cryo-EM was prepared as described above by mixing equimolar amounts of scRNA and cmpRNA, followed by precipitation in 96% EtOH+KCl, washing in 70% ice-cold EtOH, and was finally redissolved in buffer (50 mM Tris, pH 8, 100 mM MgCl_2_) to a final concentration of ∼6 mg/mL. All buffers and EtOH solutions were filtered (3 K cut-off, Amicon) prior to use. Finally, the RNA was added to grids for downstream cryo-EM analysis as described below.

### Cryo-EM data acquisition

Protochips 1.2/1.3 300 mesh Au-Flat grids were glow discharged in a GloQube Plus glow discharging system for 45 seconds at 15 mA and used immediately after for plunge freezing. Plunge freezing was performed on a Leica GP2 with the sample chamber set to 99% humidity and 15 degrees Celsius. Three microliters of sample were applied onto the foil side of the grid in the sample chamber before a 4 second delay and then 6 seconds of distance-calibrated foil-side blotting against a double layer of Whatman #1 filter paper. With no delay after blotting the sample was plunged into liquid ethane set to -184 degrees Celsius. All data were acquired at 300 keV on a Titan Krios G3i (Thermo Fisher Scientific) equipped with a K3 camera (Gatan/Ametek) and energy filter operated in EFTEM mode using a slit width of 20 eV. Data were collected over a defocus range of -0.8 to -2 micrometers with a targeted dose of 60 electrons per square Ångstrom. Automated data collection was performed with EPU, and the data was saved as gain normalized compressed tiff files with a calibrated pixel size of 0.647 Ångstrom per pixel.

### Single particle image processing and 3D reconstruction

2D classification was performed on the final particle stacks for each 3D class and are shown in Figures S2-6. Pictorial particle sorting workflows for each of the reconstructions described below can be found in Figures S7-11. Motion and CTF correction were performed in CS-Live, and the micrographs were curated to 6820 acceptable exposures ^37^.

**Class 1:** Further refinement of the small volume class failed to yield reconstructions where the helicity of the volume was clearly apparent. To attain a better reconstruction of the small volume class we performed 2D classification with 150 classes on the 668,921 extracted particles mentioned previously (Figure S2). Selection of the best classes led to a refined particle stack of 436781 particles. We used 3-class ab initio reconstruction with 60,000 randomly chosen particles to generate the initial volumes (Figure S7). This resulted in the medium volume class, the small volume class, and a larger junk volume class. The 114,396 particles from the small volume class were used to perform another ab initio reconstruction with 2 classes and 30,000 particles. Heterogeneous refinement of these volumes resulted in a class with a partially formed second helix and a class with a non-helical appendage. The 48,117 particles from the small volume class with the appendage were reconstructed with the option to force re-do GS split enabled to attain a GSFSC value of 9.5 Å.

**Class 2 and 3:** Using 3 of the 2D classes generated during CS-Live pre-processing we performed templated particle picking using a particle diameter input of 150 Å. This yielded 668,921 particles that were extracted with a box size of 512 pixels and Fourier cropped to 128 pixels. A 3-class ab initio reconstruction was performed using 60,000 randomly chosen particles, followed by heterogeneous refinement using all 668,921 particles. This yielded one junk class, one small volume class and one medium volume class. After homogeneous refinement and local refinement, the medium volume class reached a GSFSC of 6.4 Å with 243,227 particles. The medium volume class was further refined leading to class 2 and 3.

To achieve class 2, we took the 243,227 particles from the initial ab initio 3D sorting and performed a second round of classification in 3D by 5-class ab initio reconstruction followed by heterogeneous refinement (Figure S8). This resulted in a lariat class with 48,761 particles that refined to a global GSFSC of 7.7 Å. 2D classes are shown in Figure S3.

To achieve class 3, a two-class ab initio reconstruction using all 243,227 particles was performed, followed by a heterogeneous refinement that split the particles in two slightly different classes (Figure S9). One of these two classes reached a GSFSC exceeding that of the previous reconstruction with all 243,227 particles. Particles from the better class were re-extracted with re-centering using aligned shifts and a box size of 432 pixels, Fourier cropped to 216 pixels, resulting in 116769 particles. 2D classes are shown in Figure S4. Once reconstructed using the re-extracted particles we achieved a GSFSC of 6.1 Å. At this point we applied C2 symmetry to a final Local Refinement to reach a GSFSC of 5.3 Å.

**Class 4 and 5:** The particles from 2D classes that were clearly bigger than the extraction box size of 512 pixels were re-extracted with a box size of 1024 pixels. An initial volume for these larger particles was generated by 2-class ab initio reconstruction followed by heterogeneous refinement. These 33,377 particles were 2D classified again into 30 classes. The 3 best classes from this 2D classification were used for templated particle picking with an input particle diameter of 210 Å. This resulted in 202,978 particles being extracted with a box size of 1024 pixels, Fourier cropped to 256 pixels (Figure S10 and S11). These particles were sorted by heterogeneous refinement in 3D using the initial volume generated from the larger particles, a volume from the medium sized reconstructions and a volume from the smallest volume reconstruction. 130,660 particles were retained in the largest volume class while the remaining particles sorted into two junk classes.

These 130,660 particles were used for 5-class ab initio and heterogeneous refinement resulting in 1 junk class, 1 small class containing only one helix and 3 filament classes with two helices of different lengths approximating 2, 2.5 and 3 copies of the medium volume class (Figure S10). The shortest filament class contained 26,859 particles, these were used for homogeneous refinement, resulting in a GSFSC of 9.7 Å.

To isolate the largest filament particles a 3-class ab initio reconstruction followed by heterogeneous refinement of the 130,660-particle stack was performed, resulting in 1 large filament, 1 small filament and 1 junk class (Figure S11). The large filament class was used to start another 3-class ab initio reconstruction followed by heterogeneous refinement. The best class of these three volumes contained 18173 particles and achieved a GSFSC of 9.7 Å after local refinement.

### Model building

From the reconstructed volume of the medium sized particles, it was clear that the length was the correct size for a fully hybridized 36 nucleotide component and that the full volume must be comprised of two circular and two linear components. Due to the repetitive nature of the sequences, there were multiple possibilities for the hybridization of two circles with two linear complements. Different lengths of double stranded RNA were generated using RNAbuild ^36^ and fit into volume to determine how many base pairs were on each half of the circular components before the crossover. It was determined that the optimal solution was to have the short side with 14 base pairs and the longer side with 22 base pairs.

Model building was performed in UCSF ChimeraX V 1.3 ^37–40^ using ISOLDE to perform molecular dynamics with flexible fitting with a low (500 kJ mol^-1^ (mapunits)^-1^ Å^3^) weight on the map and 0 as the temperature factor ^37,39,40^. The pdb was sequence corrected and each chain renumbered using PDB-Tools ^41^. The model was then refined in real space using the Phenix software package to optimize bond angles and then relaxed in ISOLDE MDFF to reduce the clash score before a final validation using the Phenix software package ^42–46^.

### Ribozyme (TPR) catalyzed polymerization of cmpRNA

To investigate if dimer could work as a template for TPR, scRNA and cmpRNA were mixed and incubated as described above. After incubation, dimer was either purified or directly used for ribozyme polymerization assay.

For gel purification of dimer hot labeled cmpRNA and scRNA were prepared as described above and gel extracted by native PAGE. All three visible bands (band I-III in Figure 1B) were dissected from the gel and the RNA extracted while keeping the RNA at non-denaturing conditions at all times. Gel purified native RNA (normalized according to radioactive counts) was then incubated with or without active TPR (5TU/t1) (5 pmol), triplets (pppGAA and pppCUG (100 and 50 pmol, respectively)) in the reaction buffer (10 μL total volume) and frozen on dry ice followed by incubation at -7 °C for 7 days. Some samples were diluted by adding 490 uL H_2_O prior to freezing. As a positive control TPR and triplets were mixed with a linear template (same sequence as scRNA) and a fluorophore labeled primer (template: D and primer: P9, ref.: ^29^) using the same reaction mix and reaction buffer. For directly used dimer, fluorophore labeled cmpRNA (5 pmol) and scRNA (5 pmol) were prepared as described above to assemble the dimer. After dimer assembly, active TPR (5TU/t1) (5 pmol), triplets (pppGAA (100 pmol), pppCUG (50 pmol), pppGAC (50 pmol), pppAUC (50 pmol) and pppCGG (50 pmol)) and reaction buffer components were added (final volume of 10 μL in reaction buffer). For some samplens (noted in the text) additional scRNA (5 pmol) was added to the preassembled dimer and immediately before the addition of the TPR and the rest of the reactants. Finally, all samples were frozen on dry ice followed by incubation at -7 °C.

After incubation at -7 °C for 7 days, samples were thawed, adjusted to 500 μL with H_2_O and RNA was precipitation by addition of EtOH+KCl (using glycogen carrier) to 70%. Precipitated RNA was redissolved in loading buffer (95% formamide, 25 mM EDTA and Bromphemol blue) and mixed with 1 µM non-labeled complementary RNA to ensure separation of the labeled constructs form the templates. Note that this is important to release the labeled cmpRNA from the very stable dimer complex and allow detailed PAGE analysis. Finally, the samples were analyzed by denaturing PAGE. Radioactive RNA was detected using radioactive films (Thermo Scientific) and scanned in a Typhoon scanner (Thermo Scientific) using phosphorescence mode. Fluorophore labeled RNA was scanned directly in the Typhoon scanner using fluorescence mode.

### RNA hydrolysis assay

To compare RNA stability, equal amounts of gel purified dimer (band II) or single stranded cmpRNA (band I) (measured by radioactivity) was prepared in reaction buffer and incubated at -7 °C for 7 days. Following incubation, samples were investigated by denaturing PAGE and radioactivity detection as described above. Fraction of hydrolysis (per 7 days at -7 °C) was calculated as the signal below the full length cmpRNA band divided by the total signal of the lane. Calculated fraction of hydrolysis was plotted in a bar chard using Excel (Microsoft).

### Data availability

Cryo-EM map was deposited in the Electron Microscopy Data Bank under accession number EMD-19759. Atomic coordinates have been deposited into the PDB under accession number 8S6W.

## Acknowledgements

We thank the EMBION Cryo-EM Facility at iNANO, Aarhus University, for time on the Titan microscopes, technical support, and data processing on the EM computer cluster. We thank Kelly Nguyen (MRC LMB) for helpful comments on the manuscript, and Rita Rosendahl and Claus Bus for technical assistance. The research at iNANO AU was supported by the Novo Nordisk Foundation (NNF21OC0070452) (EKSM, ELK, ESA), a Carlsberg Foundation Research Infrastructure grant (CF20-0635) (ESA), and a Lundbeck fellowship (R250-2017-1502) (ELK). The research at MRC LMB was supported by the Medical Research Council, as part of United Kingdom Research and Innovation (also known as UK Research and Innovation (UKRI)) [MC_U105178804] (PH) and a Carlsberg fellowship (CF17-0809) (ELK). For the purpose of open access, the MRC Laboratory of Molecular Biology has applied a CC BY public copyright license to any Author Accepted Manuscript version arising.

