## Supplementary information for "Roles of dimeric intermediates in RNA-catalyzed rolling circle synthesis"

**Table S1. Nucleic acid sequences**

| *Oligo name* | *Seq. (5' to 3')* | *type* | *Notes:* |
|---|---|---|---|
| **Templates:** | | | |
| **scRNA** | /5Phos/GCG UUC UUC AUC UUC UUC GAU UUC UUC CAG UUC UUC | RNA | Used as either linear or circularized. |
| **cmpRNA** | UGG AAG AAA UCG AAG AAG AUG AAG AAC GCG AAG AA /3ddC/ | RNA | When noted this sequence was 5' hot labelled. |
| **F-cmpRNA** | /56-FAM/GAA GAA CUG GAA GAA AUC GAA GAA GAU GAA GAA CGC | RNA | |
| **P9** | /56-FAM/GAA GAA CUG | RNA | |
| **33 nt circle (11-GAC)** | GATCGATCTCGCCCGCGAAATTA ATACGACTCACTATA- *GTCGTCGTCGTCGTCGTCGTCGT CGTCGTCGTC*- GGGTCGGCATGGCATC | DNA | Fill-in with HDVrt (as back (Ba) primer) and in vitro transcribe. This leads to production of the *italic sequence* as the reverse compliment RNA product with the HDV ribozyme cleaved off. Cyclic phosphate was removed with PNK. |
| **36 nt circle (12-GAC)** | GATCGATCTCGCCCGCGAAATTA ATACGACTCACTATA- *GTCGTCGTCGTCGTCGTCGTCGT CGTCGTCGTCGTC*- GGGTCGGCATGGCATC | DNA | Fill-in with HDVrt (as back (Ba) primer) and in vitro transcribe. This leads to production of the *italic sequence* as the reverse compliment RNA product with the HDV ribozyme cleaved off. Cyclic phosphate was removed with PNK. |

| | | |
|---|---|---|
| **Oligonucleotides for synthesis of triplet RNA Polymerase Ribozyme (TPR):** | | |
| **5TU (Fo fill-in)** | GGATCTTCTCGATCTAACAAAAAA GACAAATCTGCCACAAAGCTTGA GAGCATCTTCGGATGCAGAGGCG GCAGCCTTCGGTGGCGCGATAGC | DNA |

| | | | |
|---|---|---|---|
| | GCCAACGTTCTCAACTATGACACGCAA | | |
| **5TU (Ba fill-in)** | CTTCTCCCTTAGCCTACCGAAGTAGCCCAGGTCGGACCGCGAGGAGGTGGAGATGCCATGCCGACCCCATGATAAACTCCATTCAACGGAGCACGCGTTTTGCGTGTCATAGTTGAGAAC | DNA | |
| **t1 (Fo fill-in)** | GACCAATCTGCCCTCAGAGCTCGAGAACATCTTCGGATGCAGAGGAGGCAGGCTTCGGTGGCGCGATAGCGCCAACGTCCTCAACCTCCAATGCATCCCACCACATGATGATGCCTGAAG | DNA | |
| **t1 (Ba fill-in)** | CTTCTCCCTTAGCCTACCGAAGTAGCCCAGGTCGGACCGCGAGGAGGTGGAGATGCCATGCCGACCCCAAAAAACCAAGGCTCTTCAGGCATCATCATGTG | DNA | |
| **5TU (final RNA product)** | GGAUCUUCUCGAUCUAACAAAAAAGACAAAUCUGCCACAAAGCUUGAGAGCAUCUUCGGAUGCAGAGGCGGCAGCCUUCGGUGGCGCGAUAGCGCCAACGUUCUCAACUAUGACACGCAAAACGCGUGCUCCGUUGAAUGGAGUUUAUCAUG | RNA | To make: Fill-in with 5TU Fo and Ba fill-in primers, then PCR with t5T7pFo and HDVrt. |
| **t1 (final RNA product)** | GACCAAUCUGCCCUCAGAGCUCGAGAACAUCUUCGGAUGCAGAGGAGGCAGGCUUCGGUGGCGCGAUAGCGCCAACGUCCUCAACCUCCAAUGCAUCCCACCACAUGAUGAUGCCUGAAGAGCCUUGGUUUUUUG | RNA | To make: Fill-in with t1 Fo and Ba fill-in primers, then PCR with t1T7pFo and HDVrt. |

| **Primers:** | | |
|---|---|---|
| **5T7 (Fo)** | GATCGATCTCGCCCGCGAAATTAATACGACTCACTATA | DNA |
| **HDVrt (Ba)** | CTTCTCCCTTAGCCTACCGAAGTAGCCCAGGTCGGACCGCGAGGAGGTGGAGATGCCATGCCGACCC | DNA |

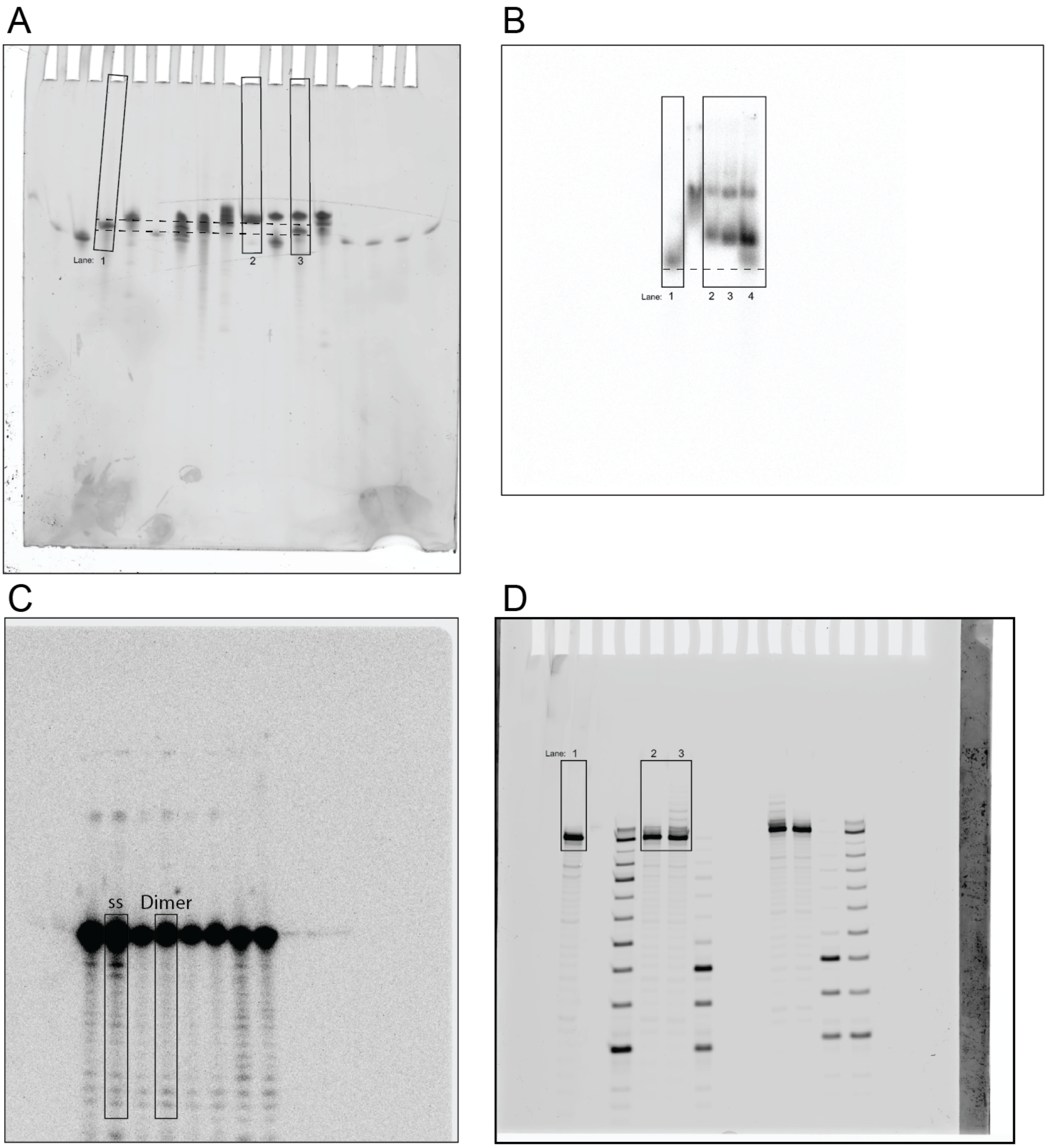

**Figure S1*.* Raw gel images. Boxes indicate regions used in main text figures.**

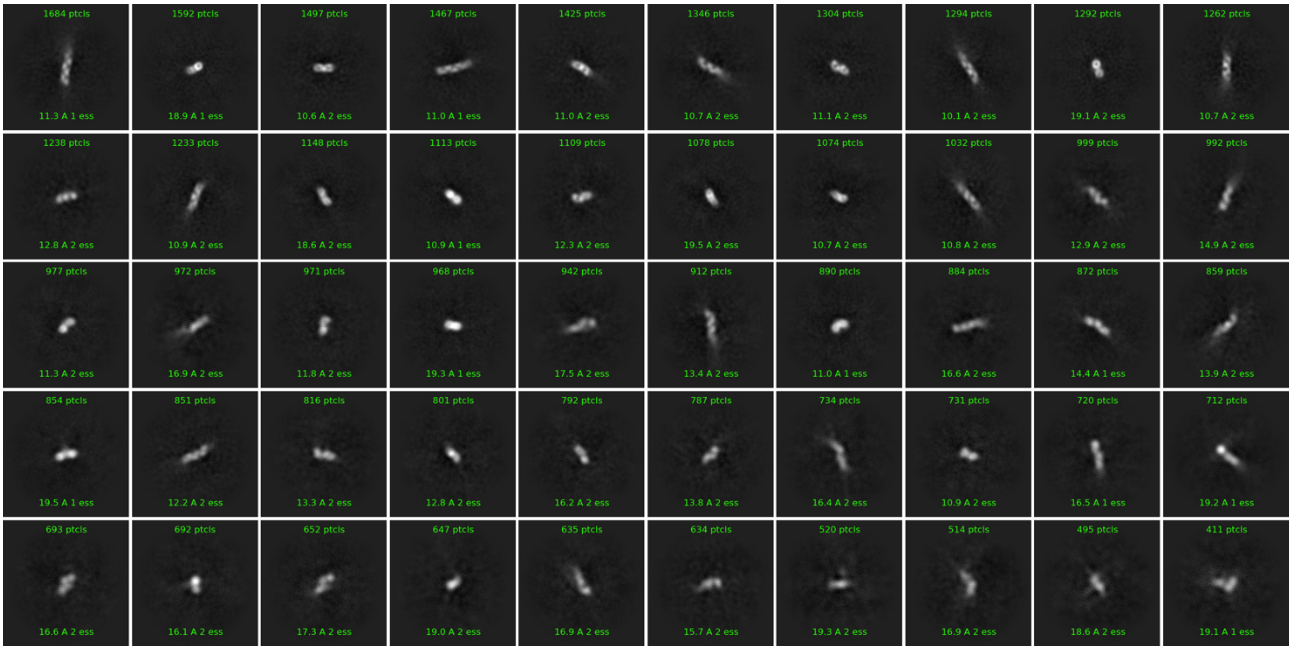

**Figure S2. Cryo-EM 2D class averages for monomer (class 1).**

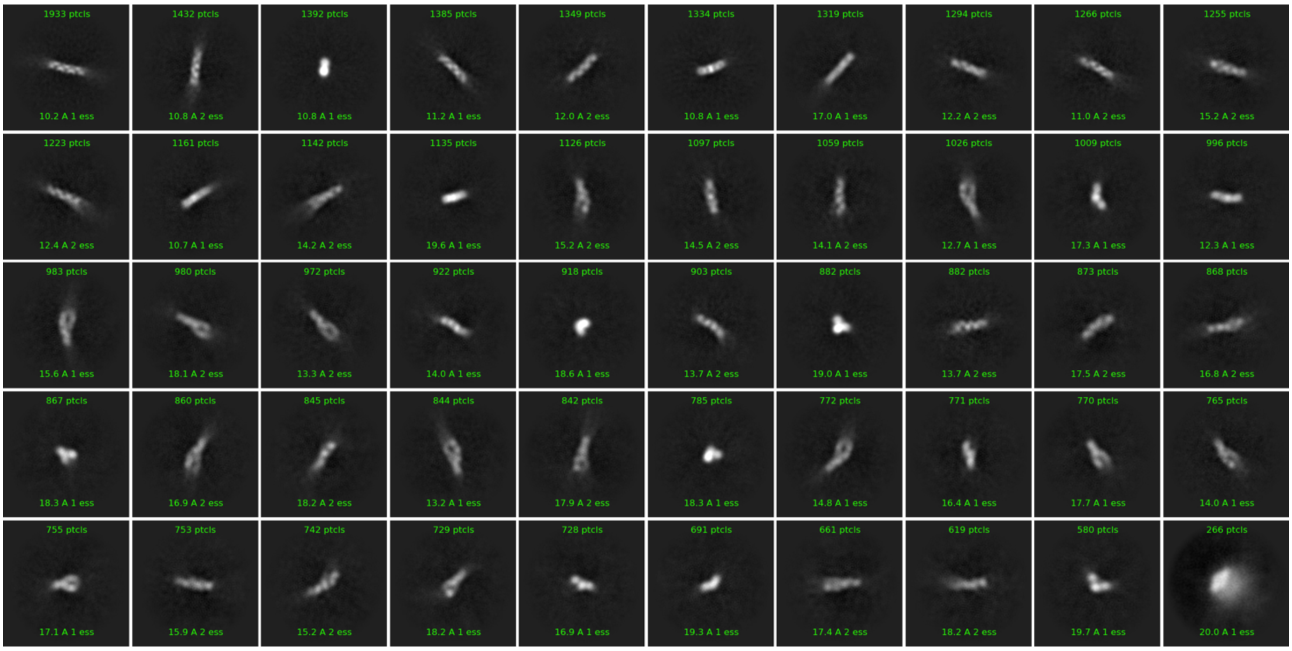

**Figure S3. Cryo-EM 2D class averages for intermediates (class 2).**

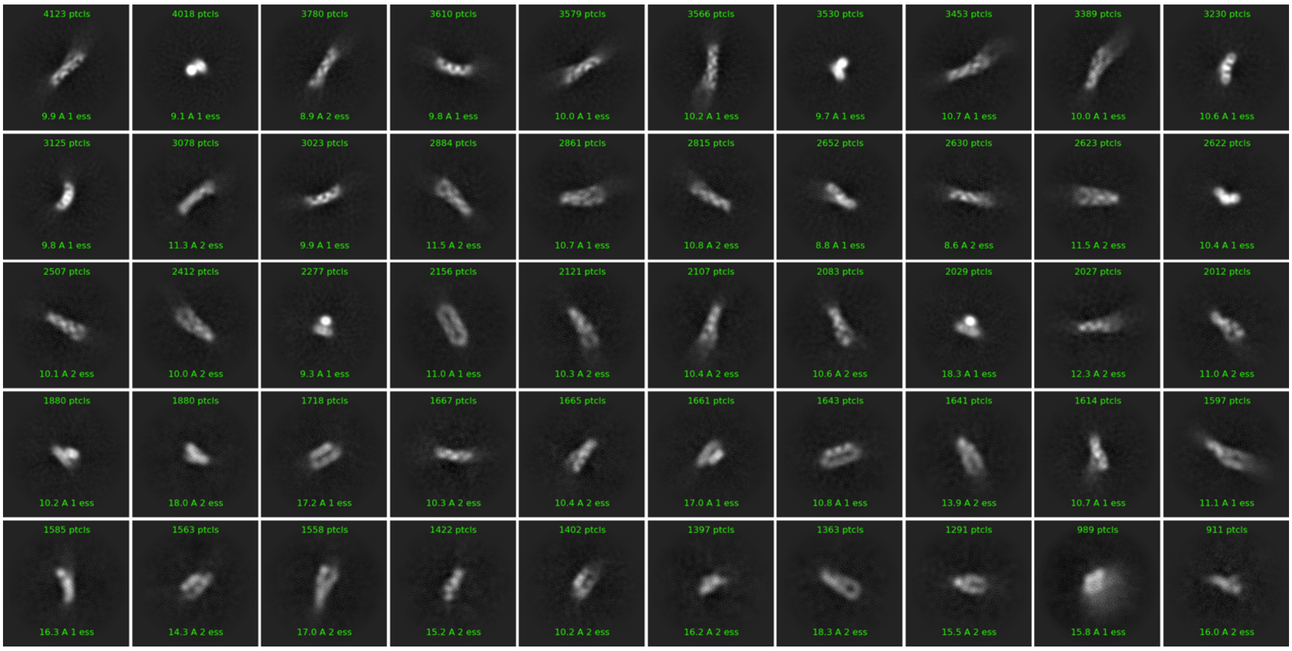

**Figure S4. Cryo-EM 2D class averages for dimer (class 3).**

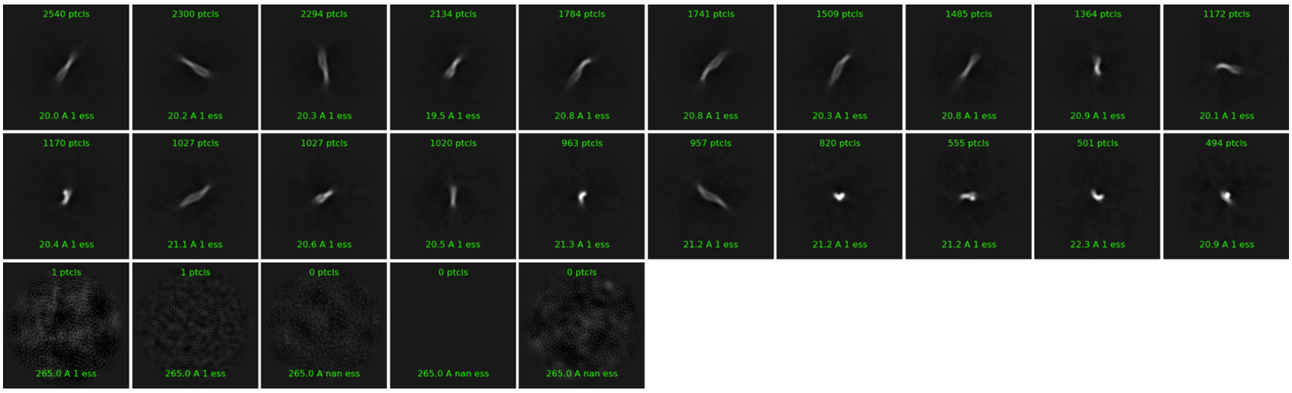

**Figure S5. Cryo-EM 2D class averages for tetramer (class 4).**

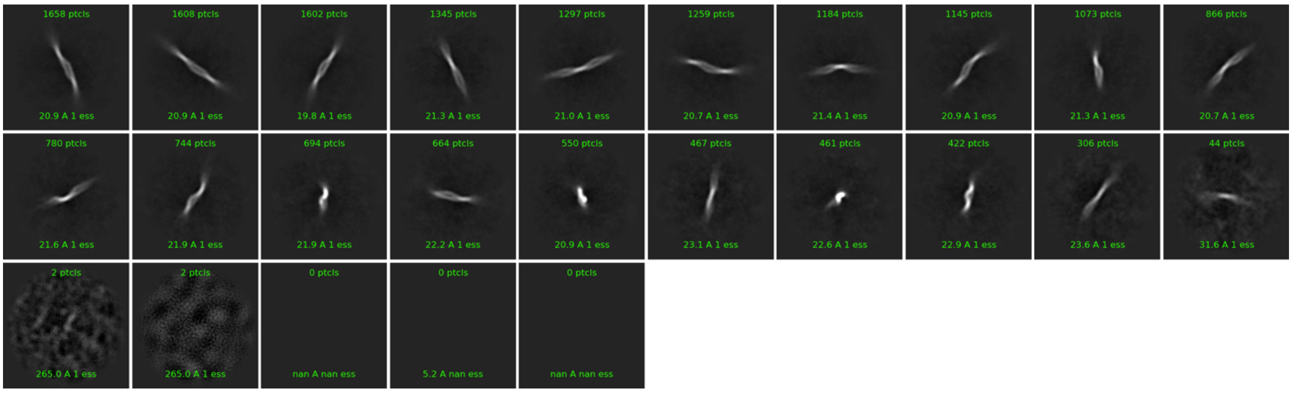

**Figure S6. Cryo-EM 2D class averages for hexamer (class 5).**

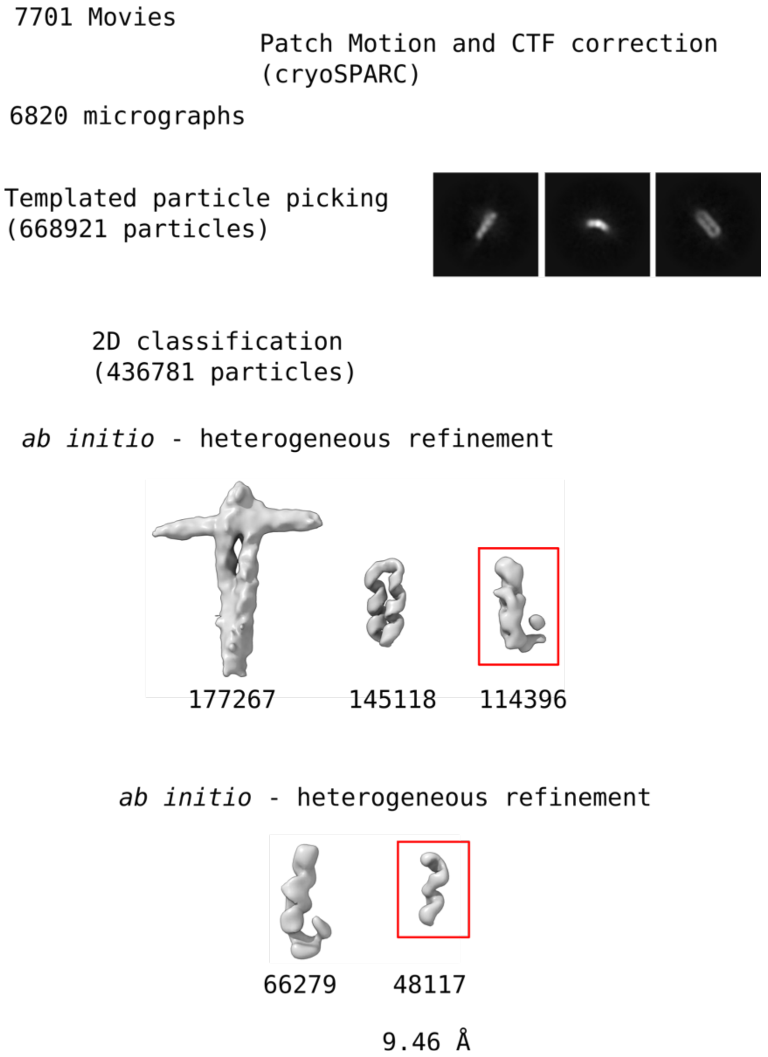

**Figure S7. Cryo-EM workflow for monomer species (class 1).**

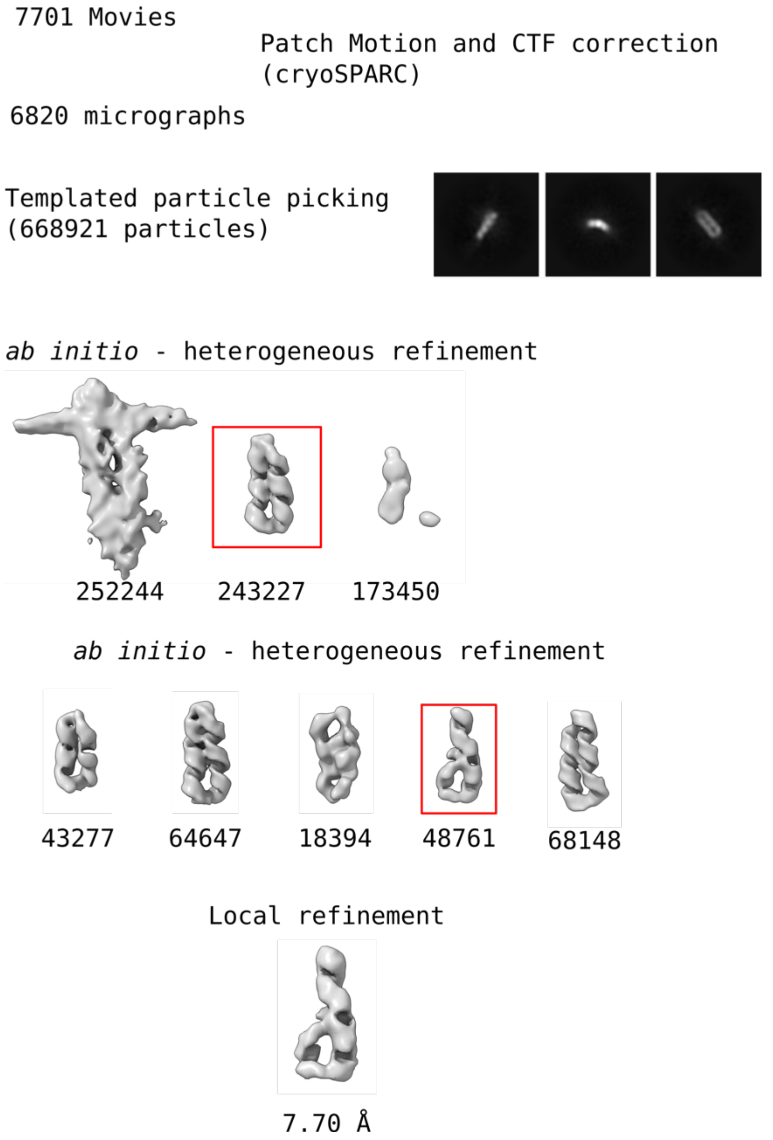

**Figure S8. Cryo-EM workflow for intermediate (class 2).**

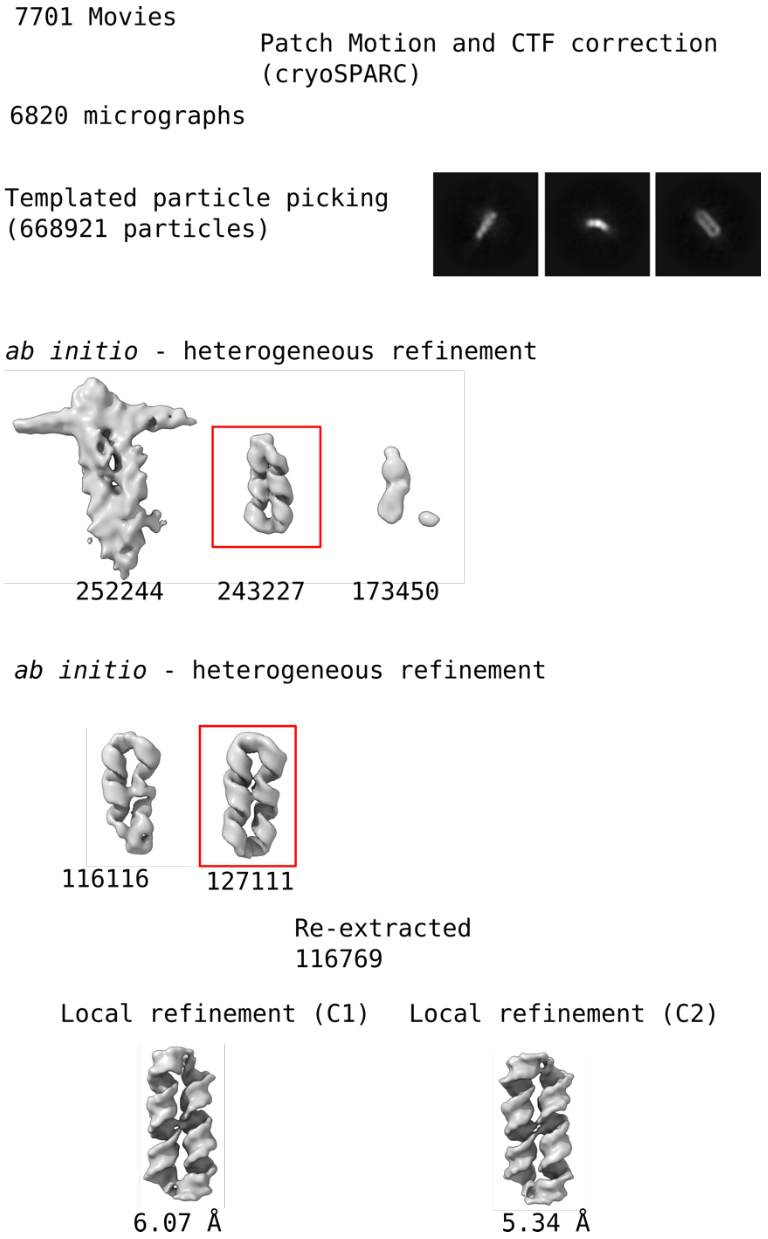

**Figure S9. Cryo-EM workflow for dimer (class 3).**

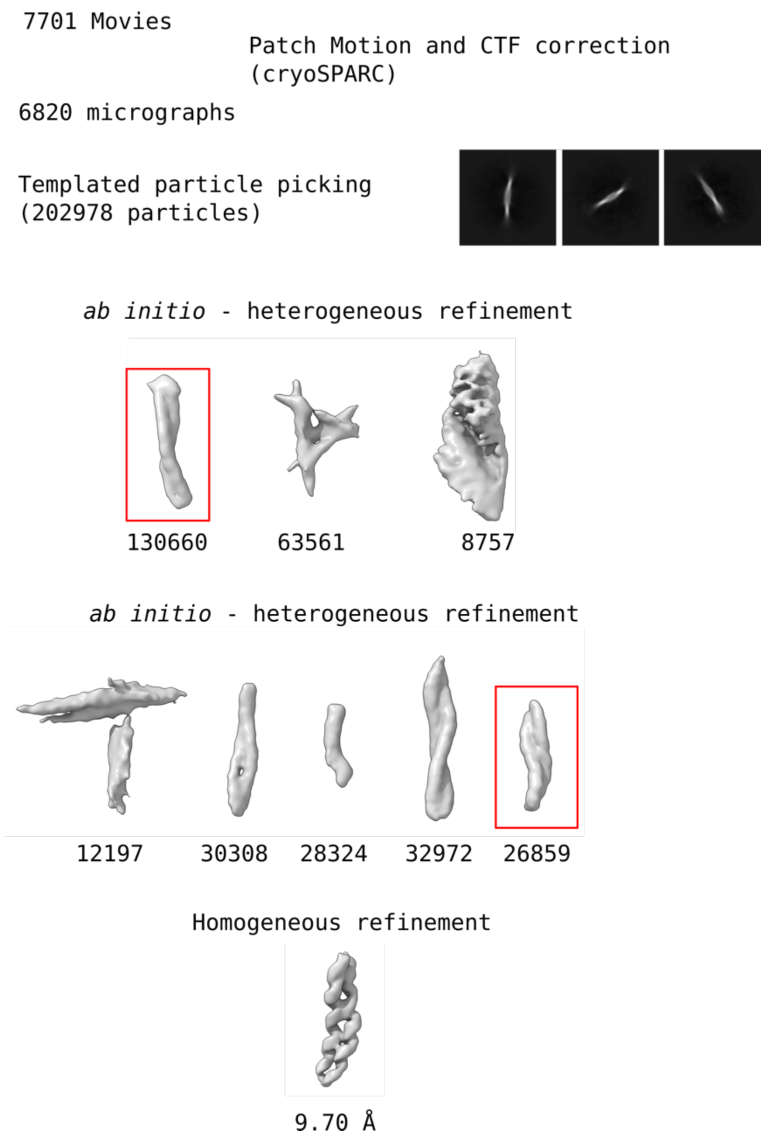

**Figure S10. Cryo-EM workflow for tetramer (class 4).**

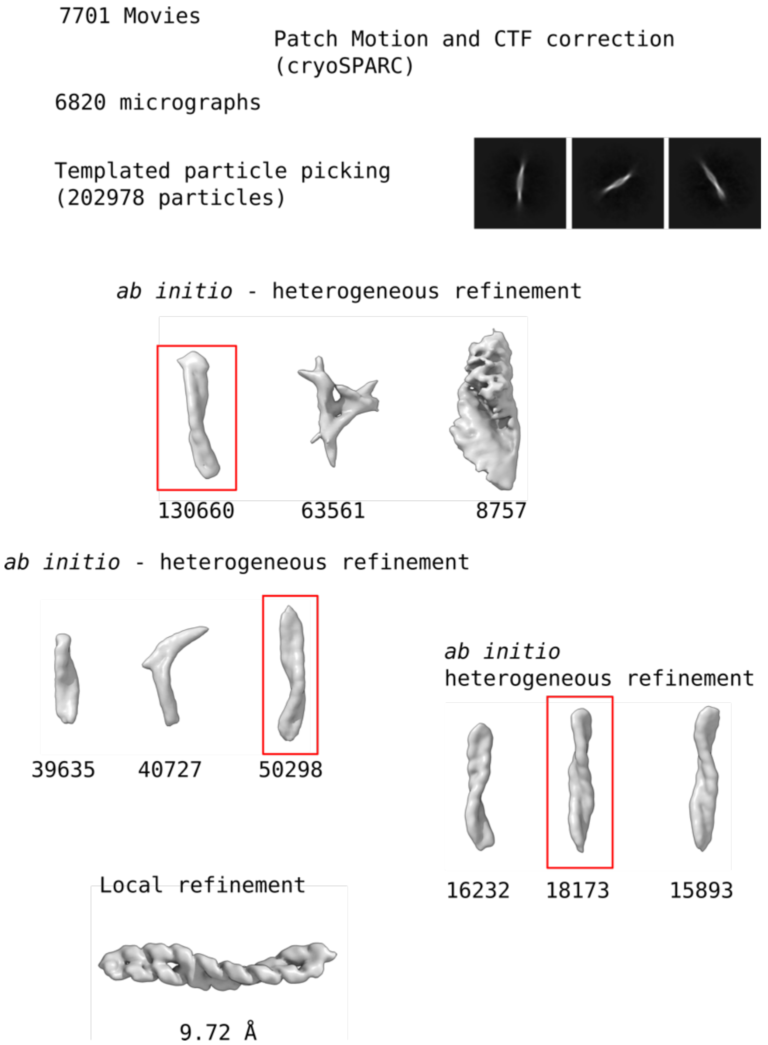

**Figure S11. Cryo-EM workflow for hexamer (class 5).**

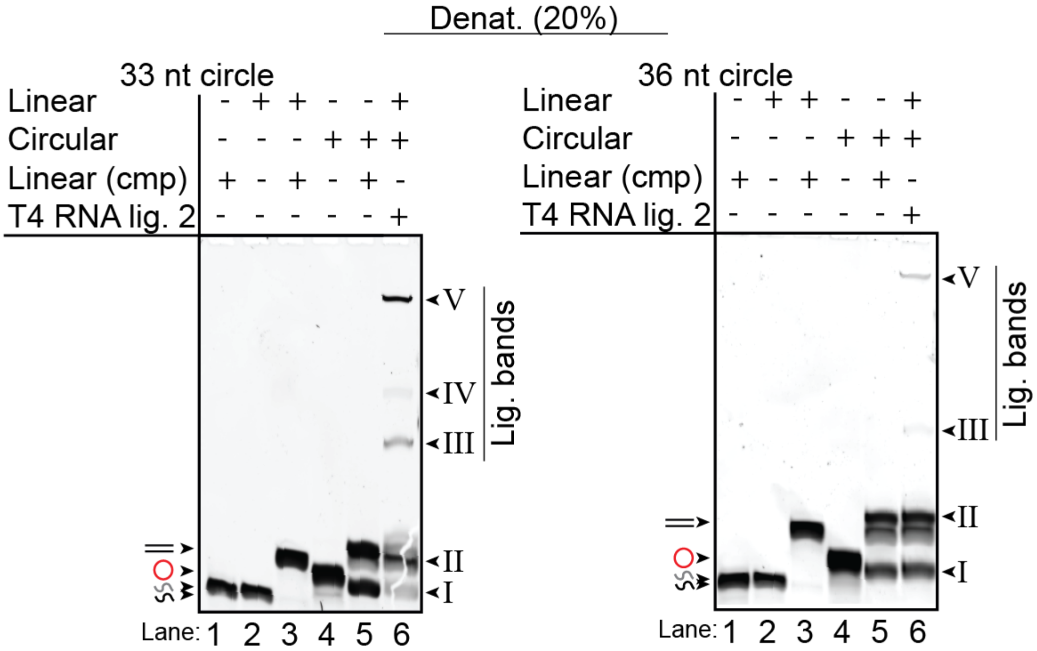

**Figure S12. Gel analysis of scRNA length, assembly, and ligation.**

Denaturing (20%, 19:1) gels of RNA strands with varying sequence and length (SYBR Gold stained). Only the stable conformations are stable under the denaturing conditions. Upon incubation of linear cmpRNA with the circular template (lane 5) stable complexes were observed (band II) similar to gels shown in main text Figure 1. Upon ligation with T4 RNA ligase 2 (lane 6) additional bands appear (Band III - V) likely corresponds to large ligation products (the RNA multimer complexes) that are now ligated together and thus stable in the denaturing gel. Interestingly, the 33 nt circles, representing a whole integer of the helical pitch (11 bp/turn), leads to much stronger ligation products compared to when the 36 nt circle, which is not a whole integer of the helical pitch.

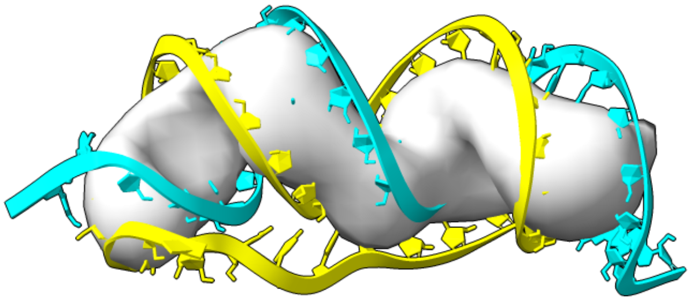

**Figure S13. Fitting of MD model to class 1 density.**

Fitting of previously modelled RNA system (in main text Ref.: 29) (consisting of 36 nt scRNA and 30 nt cmpRNA) into class 1 density.

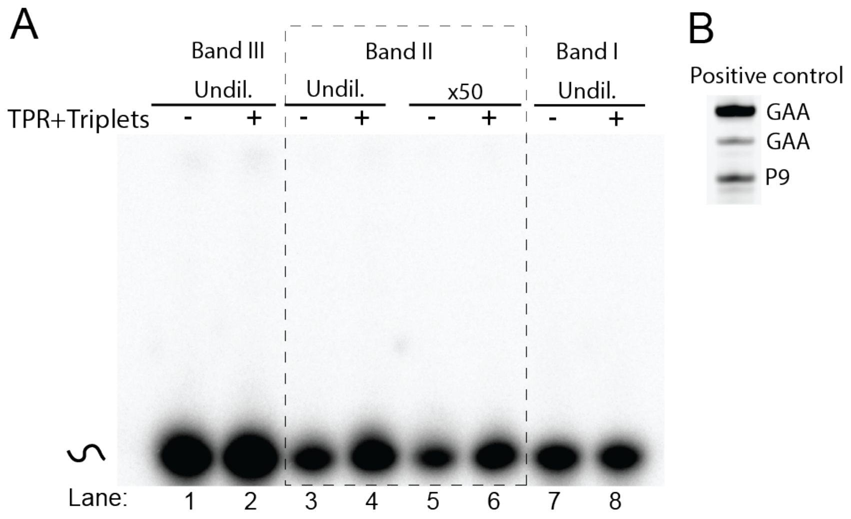

**Figure S14. Primer extension scRNA species by TPR.**

Denaturing gel (20% 19:1) of gel purified RNA after incubation with active TPR and reaction components. (A) All three bands (I-III) observed in the native gel in main text Figure 1C was gel purified and the products incubated with active TPR and triplets to extend the hot labeled cmpRNA. None of the gel purified products (the single stranded cmpRNA without template (band I, lane 1 and 2), the homodimer (band II, lane 3-6) or the multimer (band III, lane 7 and 8)) lead to extention of the cmpRNA during the reaction. x50 in lane 5 and 6 denotes that the sample had been 50-fold diluted prior to incubation, which had previously been shown to improve the RCS reaction. This did not work either. (B) Positive control showed efficient primer extension by TPR confirming that the reaction components were active.

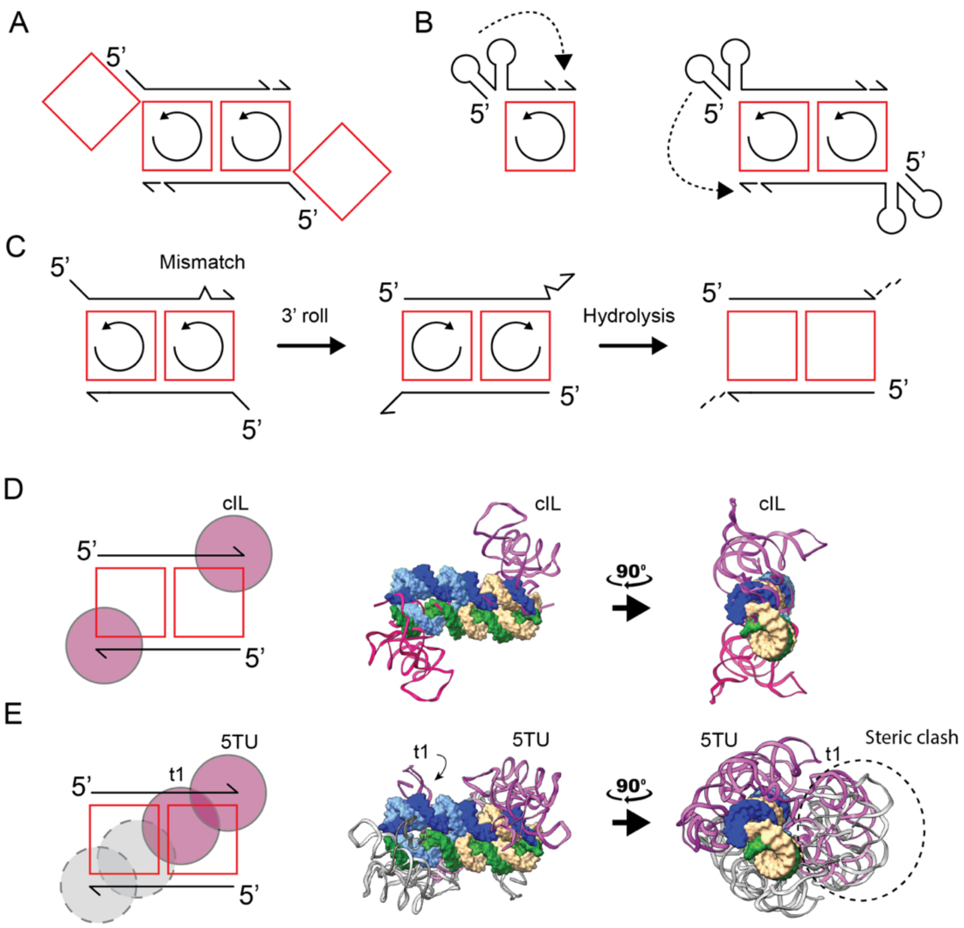

**Figure S15. Dimeric rolling circle replication hypothesis.**

(A) Schematics showing how additional circular templates may bind to 5'-end and drive directionality of synthesis. (B) Schematics of monomer and dimer RCS, where secondary structure formation drives directionality of synthesis. Dashed line with arrow indicates how a nascently folded RNA polymerase ribozyme may act within the template-product complex. (C) Schematics showing how a mismatch may induce 3' rolling and how the exposed strand may hydrolyze to reset the synthesis. (D) Model docking showing how two class I ligase (cIL) ribozymes can dock onto the dimer with the active sites positioned to ligate a triplet substrate. On the left is a cartoon illustration of the dimer (red and black) as well as the cIL (purple circles), and on the right is the full atomistic model with the dimer (shown as surface and colored as in previous figures) and the cIL (shown as cartoon and colored purple). (E) is the same as panel D but with TPR instead of cIL. Here it becomes clear that the t1 subunit of TPR collides if two ribozymes were to bind simultaneously to the dimer. Clashing regions are marked by the dashed circle. One of the TPRs is shown in purple and the other one shown in grey.
